# Deep Learning-based Identification of Intraocular Pressure-Associated Genes Influencing Trabecular Meshwork Cell and Organelle Morphology

**DOI:** 10.1101/2023.02.01.526555

**Authors:** Connor J Greatbatch, Qinyi Lu, Sandy Hung, Son N Tran, Kristof Wing, Helena Liang, Xikun Han, Tiger Zhou, Owen M Siggs, David A Mackey, Guei-Sheung Liu, Anthony L Cook, Joseph E Powell, Jamie E Craig, Stuart MacGregor, Alex W Hewitt

## Abstract

**PURPOSE:** The exact pathogenesis of primary open-angle glaucoma (POAG) is poorly understood. Genome-wide association studies (GWAS) have recently uncovered many loci associated with variation in intraocular pressure (IOP); a crucial risk factor for POAG. Artificial intelligence (AI) can be used to interrogate the effect of specific genetic knockouts on the morphology of trabecular meshwork cells (TMCs), the regulatory cells of IOP.

**METHODS:** Sixty-two genes at fifty-five loci associated with IOP variation were knocked out in primary TMC lines. All cells underwent high-throughput microscopy imaging after being stained with a five-channel fluorescent cell staining protocol. A convolutional neural network (CNN) was trained to distinguish between gene knockout and normal control cell images. The area under the receiver operator curve (AUC) metric was used to quantify morphological variation in gene knockouts to identify potential pathological perturbations.

**RESULTS:** Cells where *RALGPS1* had been perturbed demonstrated the greatest morphological variation from normal TMCs (AUC 0.851, SD 0.030), followed by *LTBP2* (AUC 0.846, SD 0.029) and *BCAS3* (AUC 0.845, SD 0.020). Of seven multi-gene loci, five had statistically significant differences in AUC (p<0.05) between genes, allowing for pathological gene prioritisation. The mitochondrial channel most frequently showed the greatest degree of morphological variation (33.9% of cell lines).

**CONCLUSIONS:** We demonstrate a robust method for functionally interrogating genome-wide association signals using high-throughput microscopy and AI. Genetic variations inducing marked morphological variation can be readily identified, allowing for the gene-based dissection of loci associated with complex traits.

## INTRODUCTION

Primary open-angle glaucoma (POAG) is a blinding disease characterised by progressive degeneration of the optic nerve and retinal nerve fibre layer.^1,2^ POAG is one of the leading causes of blindness globally.^3^ Whilst the precise pathophysiology of glaucoma is unknown, the most important modifiable risk factor is raised intraocular pressure (IOP).^1,4^ Raised IOP in POAG is primarily caused by dysfunctional aqueous humour drainage through the trabecular meshwork.^1^ Family heritage studies and genome-wide association studies (GWAS) have demonstrated a genetic contribution to trabecular meshwork dysfunction in POAG; however, the exact cellular and genetic processes involved remain unknown.^1^ Current treatments for POAG focus on reducing IOP by decreasing the production of aqueous humour or increasing outflow, with medications, or through the use of pressure-lowering surgery. However, there is currently no definitive cure for all patients with POAG.^5^ For novel pressure-lowering treatments to be developed, the pathophysiology of raised IOP in POAG must be understood, and molecular pathways for this vision-threatening disease uncovered.

Previous research has implicated a number of genes that contribute to POAG development and variation in IOP.^1,6^ Linkage analysis identified variants in the *MYOC* gene as being strongly associated with POAG.^7–9^ Disease-causing mutations in this gene have been shown to cause accumulation of a misfolded protein (myocilin), resulting in endoplasmic reticulum stress in trabecular meshwork cells (TMCs) and a subsequent rise in IOP.^6^ GWAS have identified numerous genetic variants associated with raised IOP, many of which have also been associated with POAG.^10,11^ However, further investigation into these genetic variants is required to identify which individual genes may be affected by these variants and, thus, what cellular mechanisms may be involved. The ongoing development of artificial intelligence (AI) and deep- learning tools such as convolutional neural networks (CNNs) provides a unique opportunity to investigate the genes of interest highlighted in GWAS and their effect on single cell morphology.

Deep learning is a rapidly advancing field of machine learning that relies on neural networks to learn abstract representations of data. A CNN is a specialised deep-learning model designed to learn features of image data. In supervised learning, the original images are labelled, allowing CNNs to learn the correct representation for a given label. Given the effectiveness of CNNs at image classification^12^, they have been extensively used in the analysis of cellular morphology, which is relevant in many domains of biology and medicine such as phenotype analysis,^13,14^ drug screening,^15,16^ and cell sorting.^17,18^

This study aimed to train a CNN to distinguish between primary TMCs that had specific genes from selected IOP-associated loci,^10,11^ knocked out using CRISPR/Cas and control TMCs transfected with non-targeting guide RNAs. The accuracy, as measured by the area under the receiver operator curve (AUC) metric, was used to quantify variation in morphological profiles between target gene knockouts and control cells. This high throughput approach uncovered genes at IOP loci, which, when perturbed, lead to marked variation in TMC morphology.

## METHODS

### Cell culture and passaging

Primary TMCs were collected from donors through the Lions Eye Donation service (Human Research Ethics Committee of the Royal Victorian Eye and Ear Hospital - reference number 13- 1151H). Cells were cultured in Dulbecco’s Minimal Essential Medium (Gibco, 11965118) with 10% Foetal Bovine Serum (Gibco, 16000044) and 0.5% antibiotic-antimycotic (Gibco, 15240- 062) (herein referred to as ‘culture medium’) at 37°C with 5% CO_2_. Cells were passaged by removing the culture medium and washing twice with Phosphate Buffered Saline (Gibco, 14190144). Trypsin 0.25% diluted in PBS (Gibco, 25200056) was then added, and the cells were incubated for 3 minutes at 37°C with 5% CO2. The trypsin was deactivated with cell culture medium, and cells were then aspirated into tubes and centrifuged at 1000 rpm for 5 minutes. The supernatant was aspirated, and the cell pellet was resuspended in culture medium before being plated at the desired ratio for ongoing culture. All TMCs were cultured in tissue culture treated polystyrene plates (Corning, 3516, 3524). Cell lines were tested for mycoplasma on a second weekly basis using the PCR Mycoplasma Test Kit (PromoKine, PK-CA91-1096)

### Cell transfection and CRISPR gene knockout

A total of 67 TMC lines were generated using a library of 124 targeting single guide RNAs (sgRNAs) (two for each target gene), together with 10 non-targeting sgRNAs as negative controls. SgRNAs were designed using GUIDES^19^ and are displayed in **Supplementary Table 1**. Following synthesis, sgRNAs were cloned into a novel construct that had previously been developed for the pooled single-cell RNA sequence profiling of primary cells (CROPseq-Guide- pEFS-SpCas9-p2a-puro; Addgene: #99248).^20^ The lentivirus was then packaged by transfecting HEK 293FT cells with pCMV delta 8.91, pMDG, and the recombinant plasmid via lipofectamine 2000. Lentivirus was chosen as the optimal viral vector due to its large size of ∼8.5kB allowing sgRNA, Cas9 and puromycin resistance genes to be packaged into one viral vector.^21^

Passage one primary TMCs were transfected with 50 μL of lentiviral plasmid and each CRISPR/Cas9/sgRNA/puromycin plasmid in an arrayed format. Individually cloned CRISPR/Cas9/sgRNA/puromycin plasmids were separately added to 450 μL of TMC cells in culture mixed with 1:100 lentiblast (OZ Bioscience, LB01500) in 24 well plates. Cell cultures were incubated for three days before 1 μg/mL puromycin selection occurred over four days. Transfected TMCs underwent standard cell passaging and were then resuspended in 100μl- 500μl DMEM depending on initial cell density. Initial cell density was qualitatively checked with brightfield microscopy before seeding. The predicted on-target editing efficiency for each sgRNA was generated for each sgRNA (**supplementary table 1**).The mRNA expression of each gene knockout can be quantified from RNA sequencing data, however, whilst CRISPR introduces indels into the targeted sequence, the transcription of mRNA for each target gene still occurs. Thus, directly editing efficiency is not able to be quantified using RNA sequencing data.

### Cell painting and imaging protocols

Cells were seeded at random in triplicates across 96-well plates at a density of 4.0 × 10^3^ cells per well using a Beckman Coulter MoFlo Astrios EQ fluorescence-activated cell sorter (FACS) to ensure an equal distribution of cells. The Cell Painting protocol as described by Bray and colleagues was then followed.^22^ Six fluorophores were used to highlight eight cellular components, which were imaged with high content microscopy taken at 20x magnification across five fluorescent channels on a Zeiss CellDiscoverer7 as outlined in **Table 1**. Images were auto-focussed using the definite focus strategy (a set focus point for each image) at 25 sites per well as shown in **Figure 1**.

**Table 1.** Cell Painting reagents, fluorescent channels and associated cellular organelles. The Cell Painting protocol was designed to allow a maximum number of cellular organelles to be visualised with minimal overlap of fluorescent channels.

| Cell Painting reagent | Fluorescent channel | Excitation filter (nm) | Emission filter (nm) | Organelles |
| --- | --- | --- | --- | --- |
| Hoechst 33342 | DAPI | 387/11 | 417-477 | Nucleus |
| Concanavalin A/Alexa Fluor 488 conjugate | EGFP | 472/30 | 503-538 | Endoplasmic reticulum |
| SYTO 14 Green Fluorescent Nucleic Acid stain | AF514 | 531/40 | 573-613 | Cytoplasmic RNA, nucleolus |
| Phalloidin/AlexaFluor 568<br>Wheat-Germ Agglutinin/AlexaFluor 555 conjugate | AF594 | 581/609 (phalloidin)<br>590/617 (WGA) | 622-662 | F-actin, golgi complex, cell membrane |
| MitoTracker Deep Red | AF647 | 628/40 | 672-712 | Mitochondria |

**Figure 1:**
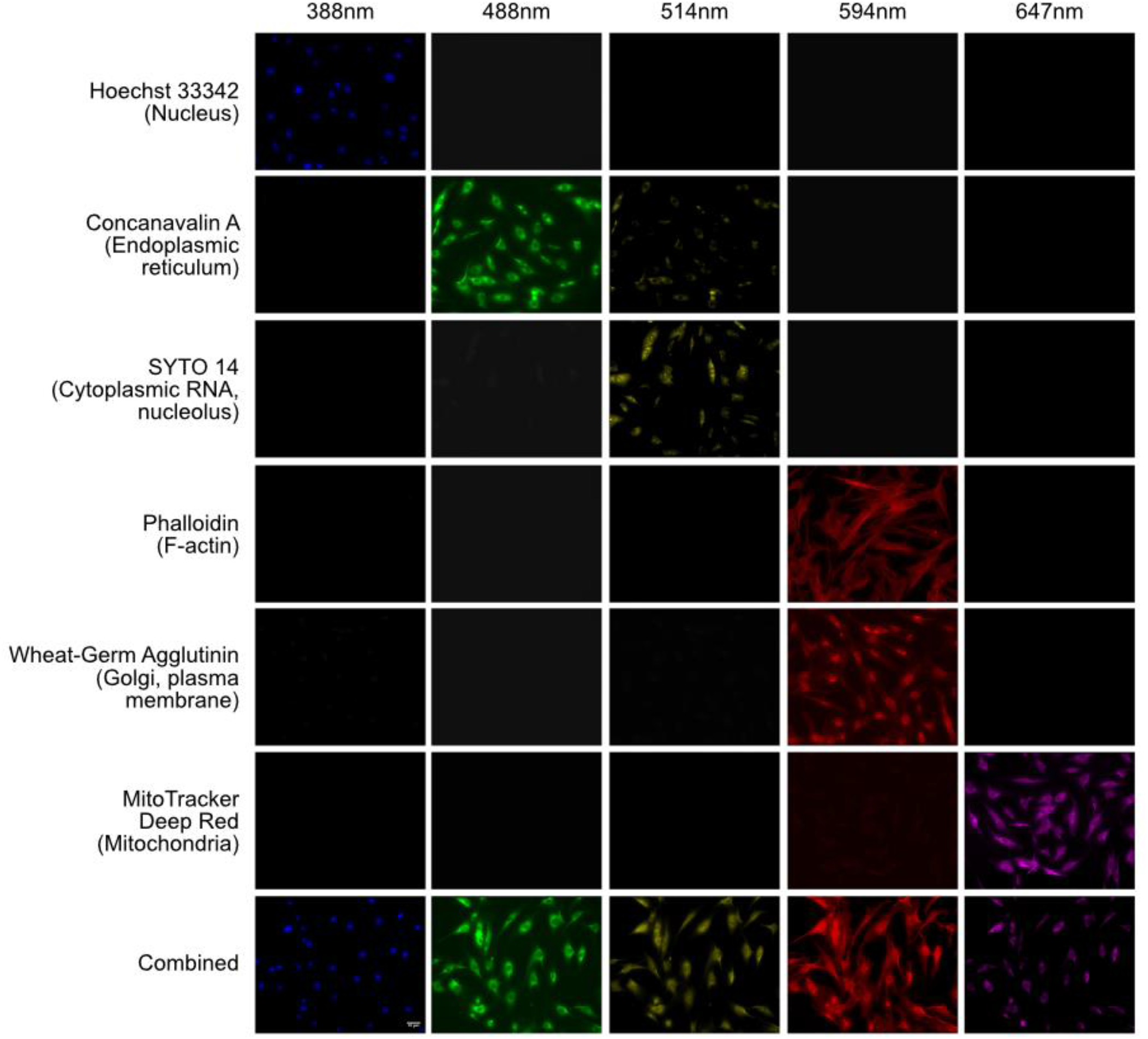
Cell Painting Assay. Example image of TMCs stained with the Cell Painting protocol in which six fluorophores are imaged over five channels to identify eight distinct intracellular organelles for morphological profiling. Each row shows cells stained with the indicated dye, or with all dyes combined (bottom row); columns indicate excitation wavelengths. Single channel testing shows minimal overlap across channels except for the Phalloidin and Wheat-Germ Agglutinin stains which are analysed together. This ensures that only a single stain will fluoresce when exposed to a particular wavelength of light. This figure shows whether a single stain would contaminate other emission channels and whether the signal of the light emission channel was dominated by the dyes we selected.

### Image preprocessing and quality control

All images were separated into multiple single-cell images using the “Save Cropped Objects” function in CellProfiler (version 3.1.9, Broad Institute, Massachusetts Institute of Technology).^23,24^ This was undertaken to ensure that single-cell morphology was the only feature of the image, and classification was not influenced by overall cell confluency. An image quality filter was then applied using CellProfiler, which flagged any low-quality images that may contain artefacts or were inadequate for analysis, and these were subsequently removed. CellProfiler analysis data was used to calculate Spearman’s rank correlation of individual cells for all cell lines. Non-correlated cells from each line were then removed by setting a Spearman correlation cutoff value of 0.15 to reduce well-to-well and batch-to-batch variation.

### CNN architecture, training and evaluation

The CNN architecture is outlined in **Supplementary Table 2** and accessible via GitHub. The dataset was first split into training (80%), validation (10%), and testing (10%) sets. A separate CNN was trained for each fluorescent channel of each gene across five replicates (each with a different random seed to create individual datasets). Training was conducted for 100 epochs, with the model being saved at each epoch. An Adam optimiser was used with a learning rate of 0.0001. For evaluation, the best-performing model of the 100 epochs as per the loss function was selected and evaluated on the test set. The AUC metric was used to quantify CNN performance and thus the degree of morphological variation induced by genetic variations. The highest-performing models were all selected prior to reaching 100 epochs where model overfitting began to reduce model accuracy.

## RESULTS

### Image Filtering and Data Split

Filtering using CellProfiler and by Spearman correlation reduced the total dataset size from 225,095 images per channel to 114,830 images per channel, yielding a total of 574,150 images for analysis. The proportion of images removed via Spearman filtering varied across groups from 22.1% (*ANTXR1*) to 70.0% (non-targeting group one). The five non-targeting control lines had the greatest proportion of images removed via Spearman filtering as shown in **Figure 2**. The total number of cell images after filtering ranged from 221 (*ADAMTS6*) to 4323 (*ANTXR1*). This inter-group variability was balanced during training with image rotation data augmentation (0, 90, 180, 270, with or without horizontal mirroring) to reach approximately 3,000 images per group. A random selection of non-targeting control images was then selected to produce a balanced dataset of gene knockout and non-targeting control images. The same non-targeting images were chosen for each knockout comparison. The dataset was split into training (80%), validation (10%), and testing (10%) sets.

**Figure 2:**
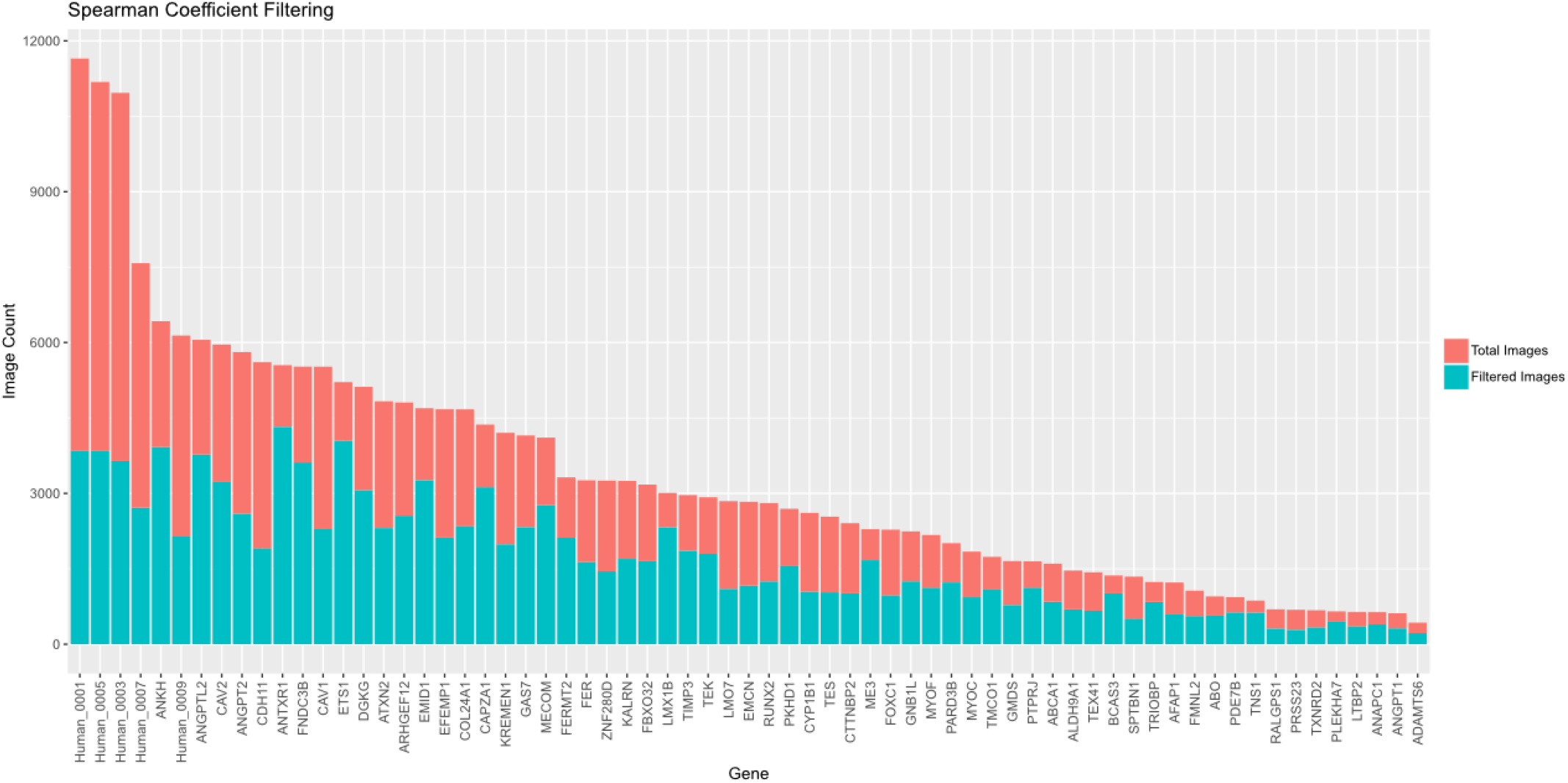
Total number of images for each arrayed cell line following Spearman correlation filtering. Images were removed from the dataset if the Spearman correlation was >0.15 in order to improve the quality of the dataset and reduce the effect of well-to-well and batch-to-batch variation. Ultimately, the percentage of cells removed ranged from 67% (control line 1) to 22% (*ANTXR1*).

### Overall morphological variation induced by genetic knockouts

The AUC metric was used to assess the ability of the CNN to distinguish genetic knockout lines from non-targeting control lines thus providing a quantifiable value of morphological variation induced by gene knockouts. The mean AUC of five replicates across five channels was calculated to produce an overall AUC for each target gene. Knockout of *RALGPS1* produced the most morphologically distinct TMCs (AUC 0.851, SD 0.030), followed by *LTBP2* (AUC 0.846, SD 0.029) and *BCAS3* (AUC 0.845, SD 0.020). The overall AUCs ranged from 0.564 (*LMO7*) to the most distinguishable at 0.851 (*RALGPS1*) as displayed in **Figure 3**.

**Figure 3:**
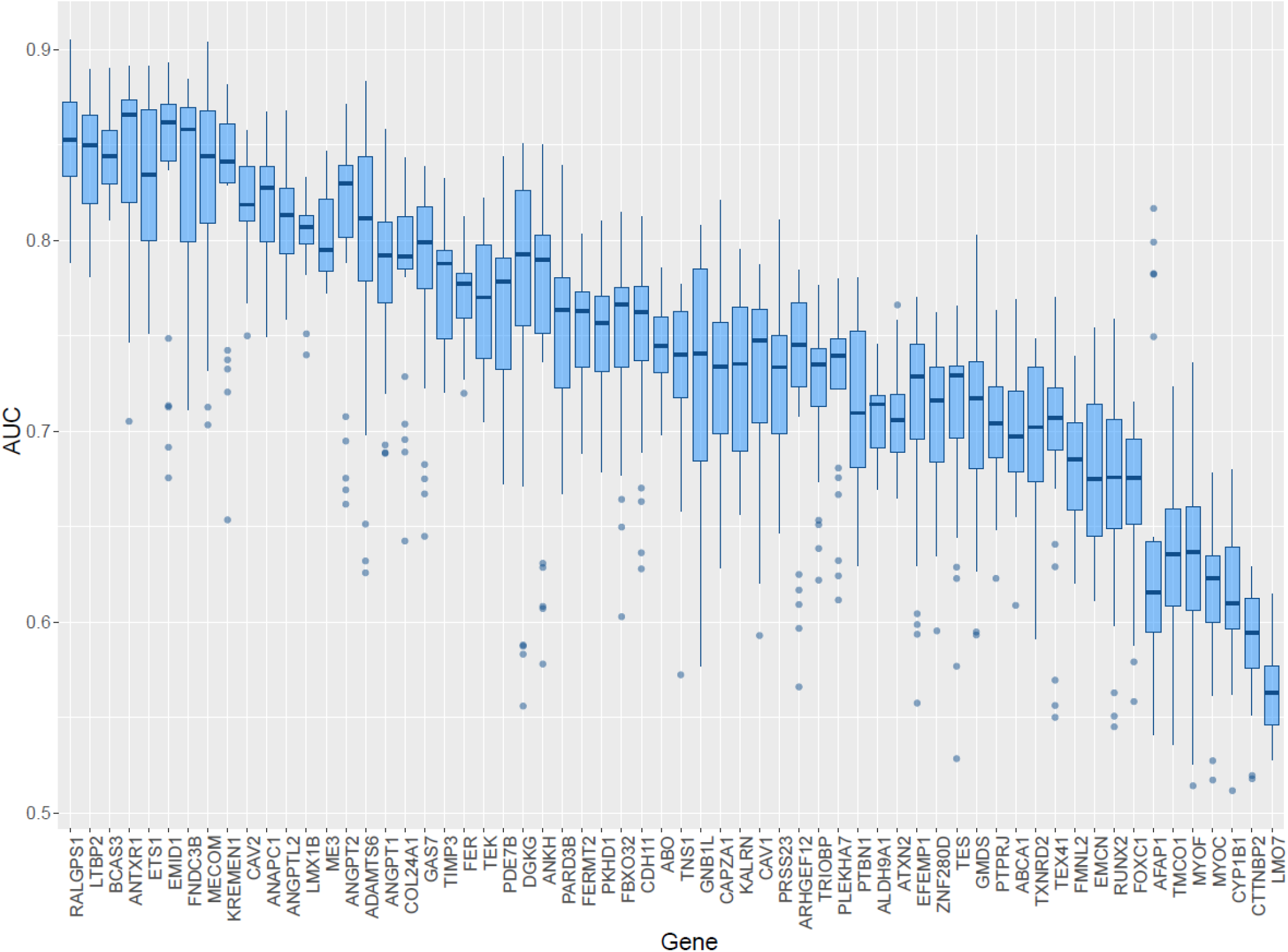
Mean CNN AUC scores for each gene-knockout cell-line. The mean AUC score when training a CNN to distinguish between gene-knockout cell-lines and non-targeting control cell-lines. A higher AUC indicates a more distinct morphological variation induced by a particular gene-knockout. The gene knockouts are ordered in decreasing order of mean AUC across all organelles. The bars represent the median AUC with upper and lower quartile boxes. Outliers are displayed as single dots.

### Morphological variation induced in individual organelles

Twenty one (33.9%) gene knockout groups had greater morphological distinction in the mitochondrial channel (mean AUC 0.760 of all cell lines, SD 0.070) compared to other organelles, illustrating that mitochondrial variation occurs in a large proportion of the gene knockouts. The relative AUC of each gene across all organelles is shown in **Figure 4**. Endoplasmic reticulum showed the next greatest morphological variation evident in 16 (25.8%) of the gene knockout lines (mean AUC 0.756, SD 0.079). The F-actin/cell membrane/Golgi body channel showed the highest morphological variation in 13 (20.9%) gene knockout lines (mean AUC 0.751, SD 0.073) followed by 11 (17.7%) knockout lines in the cytoplasmic RNA/nucleolus channels (mean AUC 0.753, SD 0.078). Finally, only the *ANAPC1* knockout showed morphological variation most in the nucleus (mean AUC 0.677, SD 0.079).

**Figure 4:**
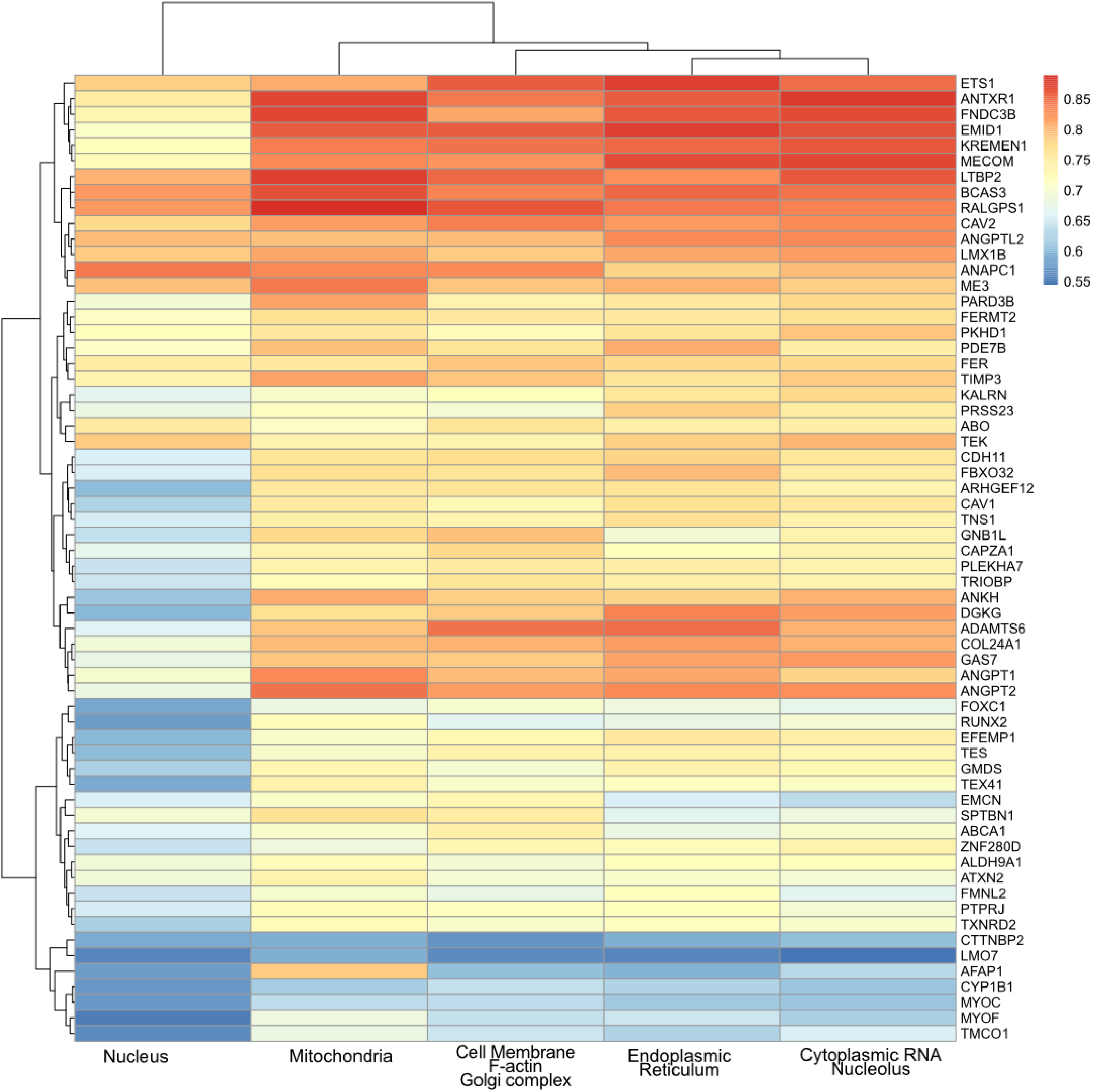
Gene knockout cell line AUC for each organelle. Heatmap of the morphological variation (AUC) across individual fluorescent channels for each gene knockout. Red shading indicates a higher degree of morphological variation as indicated by a higher AUC.

### Gene prioritisation

Finally, we used the trained CNN AUC metrics to investigate TMC morphological variation for genes at multi-gene loci.^10,11^ **Table 2** displays the AUC (knock-out of target gene compared to non-targeting control) for 15 genes across seven loci. For five of these loci, we identified gene knockouts (*ALDH9A1, CAV2, ME3, RALGPS1* (present in two loci)) which resulted in greater morphological variation than knockout of their neighbouring gene counterparts. Knockout of genes at two multi-gene loci (*EMID1-KREMEN1* and *GNB1L-TXNRD2*) generated TMCs that were morphologically similar and thus, could not be prioritised.

**Table 2:**
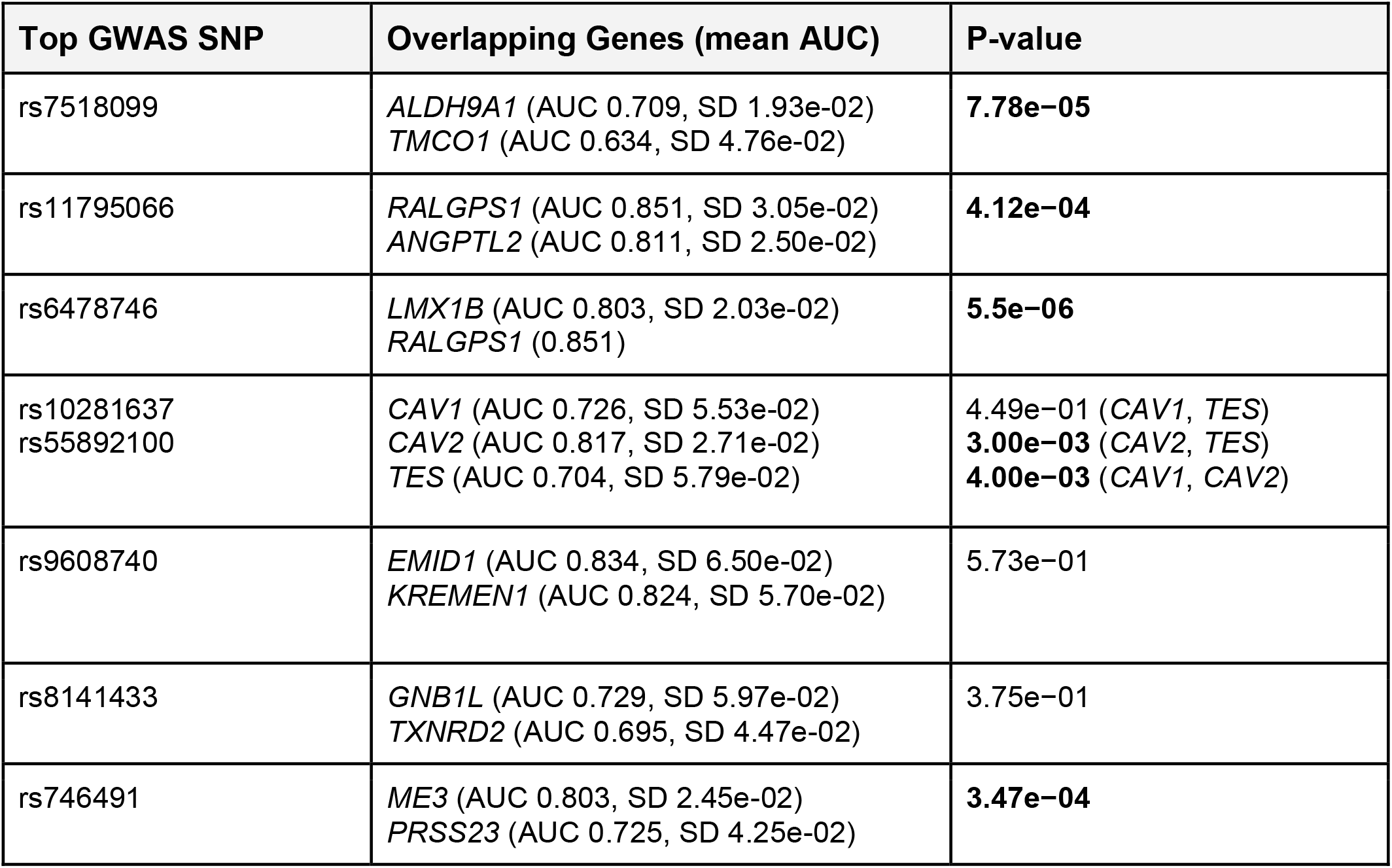
Comparison of CNNs to morphologically distinguish TMCs with knockout of genes at overlapping IOP-associated loci.^10^. The mean AUC across all fluorescent channels of target knockouts versus non-targeting control cells was compared for genes at the same locus. A higher AUC indicates a larger degree of morphological variation compared to normal control cells. This allows for prioritisation of overlapping genes at given loci.

## DISCUSSION

There has been a shift in recent years towards using high-throughput profiling to undertake large-scale studies investigating the cellular basis of disease. This shift has been accelerated by advancements in computational technology and AI as a method of rapidly analysing large, complex datasets. In this study, we utilised a convolutional neural network to perform a high-throughput morphological analysis of genetic variations associated with IOP variation in primary human TMCs. By training the CNN to distinguish gene knockout cells from healthy control cells, we could use the AUC as a metric to quantify differences in cellular morphology induced by various genetic variations. Therefore, the AUC can be used to identify which variations invoke a greater degree of morphological change and, thus, which are more likely to be involved in IOP dysregulation and the pathogenesis of POAG.

Of the genes known to cause primary congenital glaucoma or anterior segment dysgenesis, *LTBP2* and *TEK* showed marked differentiation from normal control morphology. The *LTBP2* knockout cell line was readily distinguished from normal control TMCs (AUC 0.846) with the greatest degree of difference occurring in mitochondrial morphology indicating that *LTBP2* may play a role in mitochondrial function. *LTBP2* encodes for latent transforming growth factor beta binding protein 2 which is an extracellular matrix protein associated with fibrillin-1 containing microfibrils and is hypothesised to modulate extracellular matrix production.^25^ Variations in *LTBP2* have been previously associated with primary congenital glaucoma, microspherophakia, megalocornea and Weill-Marchesani syndrome.^25–28^ A previous study has identified that *LTBP2* knockout may contribute to the development of POAG via dysregulation of the extracellular matrix; a crucial component of the trabecular meshwork.^29^ Studies looking at dilated cardiomyopathy and right ventricular failure have also implicated *LTBP2* function in fibrosis regulation which may indicate a role in the pathogenesis of trabecular meshwork dysfunction.^30,31^

The *TEK* knockout cell line also showed significant differentiation (AUC 0.768) most prominent in the cytoplasmic RNA and nucleolus channel. This gene encodes for a tyrosine-kinase receptor and is highly involved in the regulation of angiogenesis and vascular stability.^32^ It also acts as a receptor for *ANGPT1* which has been shown to be crucial for development of Schlemm’s canal.^33–35^ Variations in *TEK* have been associated with raised IOP and congenital glaucoma primarily due to disruption of Schlemm’s canal, indicating a potential interaction with *ANGPT1* in the development of glaucoma.^35–38^ Curiously, *MYOC, CYP1B1, GMDS*, and *FOXC1* knockouts resulted in only mild differentiation from control TMC morphology (AUC 0.615, 0.612, 0.704, 0.665, respectively) despite an association with glaucoma and anterior segment dysgenesis.^7,39–42^ These gene knocktous may not invoke significant morphological variation as they are primarily involved trabecular meshwork development rather than the maintenance.^43^ Furthermore, some gene mutations associated with congenital glaucoma are gain-of-function mutations and therefore will not show significant change when knocked out. Another reason for not seeing change in cellular morphology is that these genes may primarily act extracellularly such as *MYOC* which has been shown to demonstrate accumulation of extracellular products in specific mutations.^44^

The knockout of *RALGPS1* resulted in the greatest degree of differentiation (AUC 0.851) compared to other cell lines and was most prominent in the mitochondrial channel. This gene encodes for ras-specific guanine nucleotide-releasing factor RalGPS1, which is involved in Ras protein activation.^45^ Not only has *RALGPS1* been associated with raised IOP^10,11^, but previous studies have also highlighted a link to high myopia as well as a role in optic nerve regeneration.^46,47^ The *BCAS3* gene knockout also produced a high degree of differentiation (AUC 0.845), which was also greatest in the mitochondrial channel. This gene encodes for breast carcinoma-amplified sequence 3 and has been shown to play a role in angiogenesis.^48,49^ *BCAS3* variants have been previously associated with glaucoma and optic nerve head parameters.^50–52^

Overall, the mitochondrial channel most frequently displayed the greatest degree of differentiation (33.9% of all cell lines). Previous studies have highlighted an association between glaucoma and mitochondrial dysfunction, likely related to the high energy requirements of retinal ganglion cells.^53–55^ Studies have also shown direct evidence of mitochondrial dysfunction in POAG affected eyes indicated by increased mitochondrial respiratory activity and elevated retinal mitochondrial flavoprotein; both of which are associated with mitochondrial dysfunction.^56–58^ The endoplasmic reticulum channel also showed the most morphological variation in a large proportion of cell lines (25.8%), which is in keeping with many studies that have highlighted a link between glaucoma and endoplasmic reticulum stress.^59–61^ Interestingly, the *ANAPC1* knockout was the only cell line to display the greatest differentiation in the nucleus channel compared to other organelles. Similarly, this gene is involved in progression through cellular mitosis.^62^

This work introduced a novel method for prioritising genes at overlapping loci identified in GWAS using CNN analysis.^10,11^ The results show that *ALDH9A1, RALGPS1, CAV2* and *ME3* show statistically greater differentiation from control cells than the respectively associated gene at the same locus. Studies have previously associated POAG with genetic variants at the inter- genomic region of *TMCO1* and *ALDH9A1*.^63–65^ The results of this study point toward *ALDH9A1* being the implicated gene in POAG due to inducing a greater degree of morphological change compared to *TMCO1* (p-value 7.78e−05). The mitochondrial channel in *ALDH9A1* displayed the greatest degree of differentiation, highlighting the potential role of mitochondrial dysfunction in *ALDH9A1* interruption in POAG. This is supported by the role of *ALDH9A1* in carnitine synthesis, which takes place in the mitochondrial matrix.^66^ There have also been numerous studies illustrating an association between POAG and variations at the inter-genomic region of *CAV1* and *CAV2*.^67–70^ This analysis prioritised *CAV2* as a potential causative gene, with a higher degree of morphological change from control cells than *CAV1* (p-value 4.00e−03). The *CAV2* knockout cell line displayed the most prominent changes in the F-actin, Golgi complex, cell membrane fluorescent channel. Supporting this, previous studies have highlighted the interaction between *CAV2* and the Golgi complex.^71–73^ The genomic region containing *ME3* and *PRSS23* has previously been associated with open-angle glaucoma.^74^ Our study highlighted a statistically greater degree of morphological change in the *ME3* cell line providing evidence for prioritisation over *PRSS23* in the pathogenesis of POAG. The remaining genes at overlapping loci (*EMID1 vs KREMEN1* and *GNB1L vs TXNRD2*) showed no statistically significant differences in morphology. They will require further investigation to prioritise which of these may be the causative gene.

A further application of AI-based analysis of single cell morphology is to predict gene expression as demonstrated in prior studies. For example, Chlis and colleagues developed a machine learning model to predict gene expression of human mononuclear blood cells and mouse myeloid progenitor cells based on cellular morphology.^75^ Our study further highlights the complex interaction between cell morphology and gene expression and the opportunity that AI poses as a means of analysing the large amounts of data produced. Further investigation into this field could uncover the genetic drivers behind pathological changes in morphology that drive disease processes and allow for identification of novel therapeutic targets.^75,76^

One of the main limitations of this study lies in the intrinsic difficulty in interpreting the decision-making process of CNNs. This means it can be difficult to establish if morphological features learned by the CNN are truly pathological or simply due to systematic bias. For example, if wells had lower cell density, the cells may grow to a larger size, thus cell size may inadvertently influence the decision-making of the CNN.

In summary, this study used a powerful approach to quantify morphological change induced by genetic variations associated with POAG. *RALGPS1* produced the greatest morphological variation. As well as this, we could prioritise genes at overlapping loci identified to have an association with IOP. However, there are some limitations due to the difficulty in removing systematic bias from the methodology. This bias may result in the CNN learning features that are not directly associated with IOP physiology. This study highlights a new avenue for utilising CNNs trained on single-cell morphology to further interpret the results of GWAS.

## Acknowledgements

D.A.M., J.E.C., J.E.P, S.M., and A.W.H. are supported by the Australian National Health and Medical Research Council (NHMRC) Fellowships. X.H. was supported by the University of Queensland Research Training Scholarship and QIMR Berghofer Medical Research Institute PhD Top Up Scholarship. We are grateful for funding from Australian Vision Research; a NHMRC Program grant (1150144), Partnership grant (1132454) and the Clifford Craig Foundation.

## SUPPLEMENTARY

### Code availability

The Python functions utilised for data preparation, CNN training and evaluation are available on GitHub: https://github.com/ConnorG1/TMC_CNN

### Data availability

Data is available at the European Bioimage Institute Bioimage Archive: Accession S-BSST841

**Supplementary table 1.**
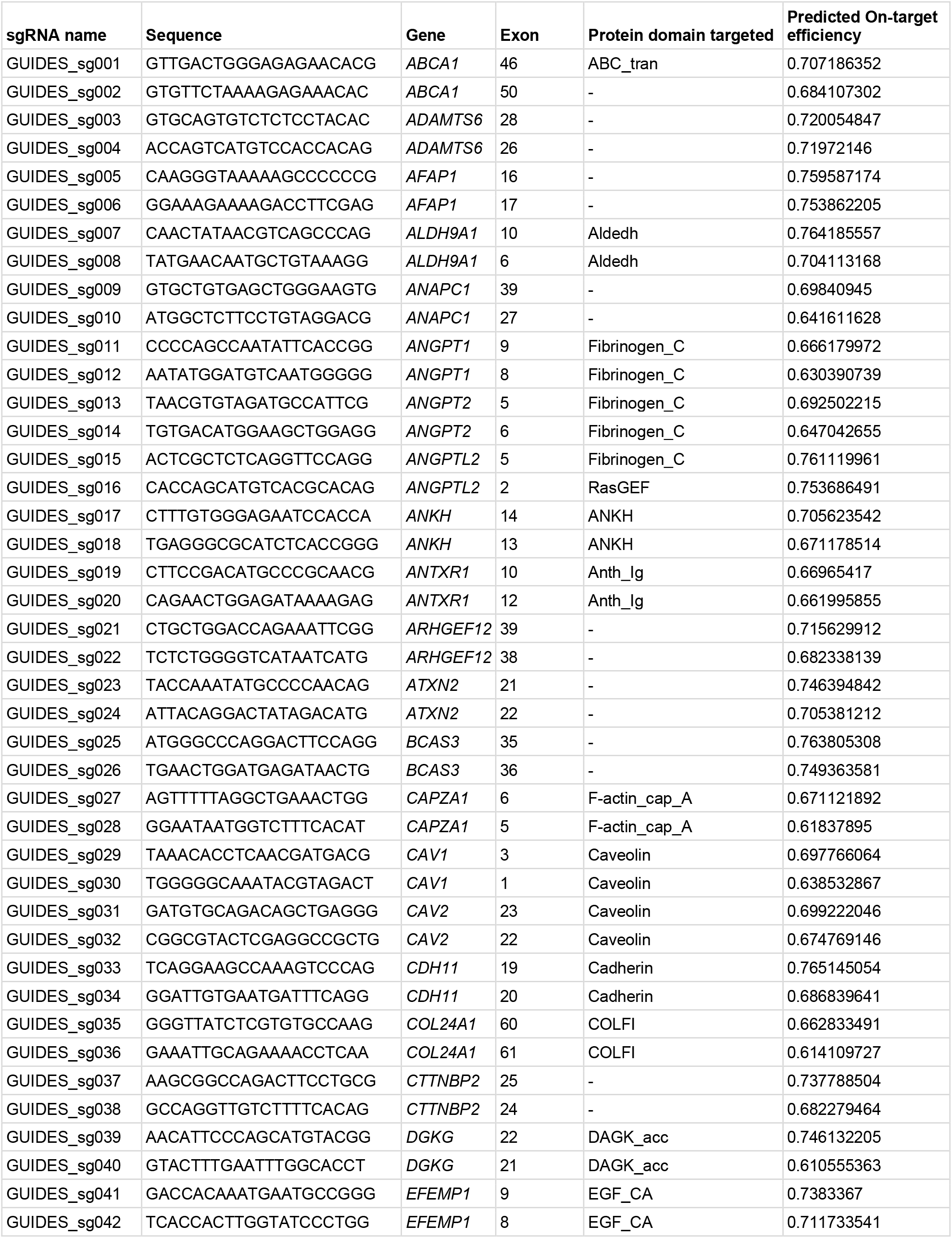
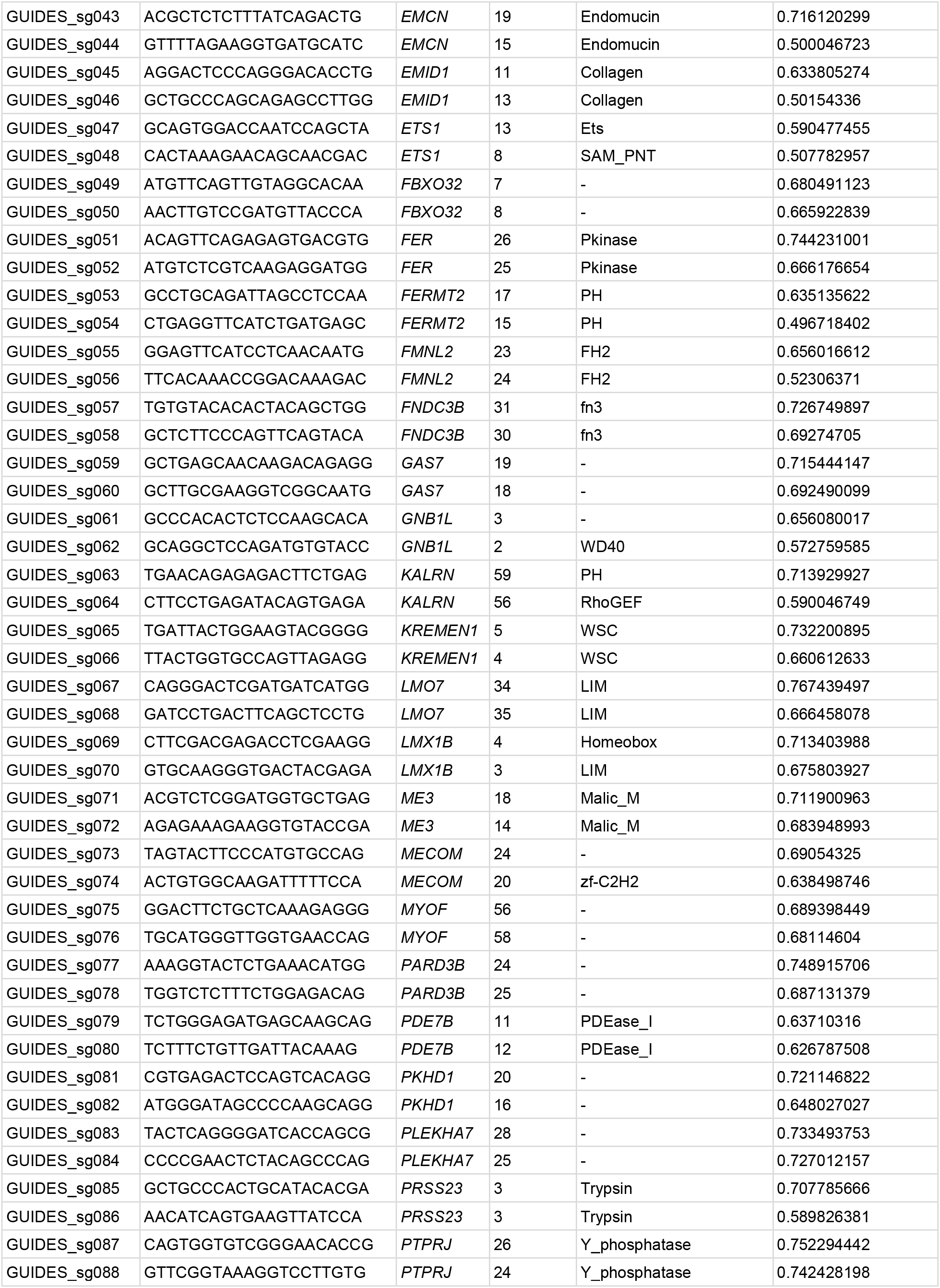
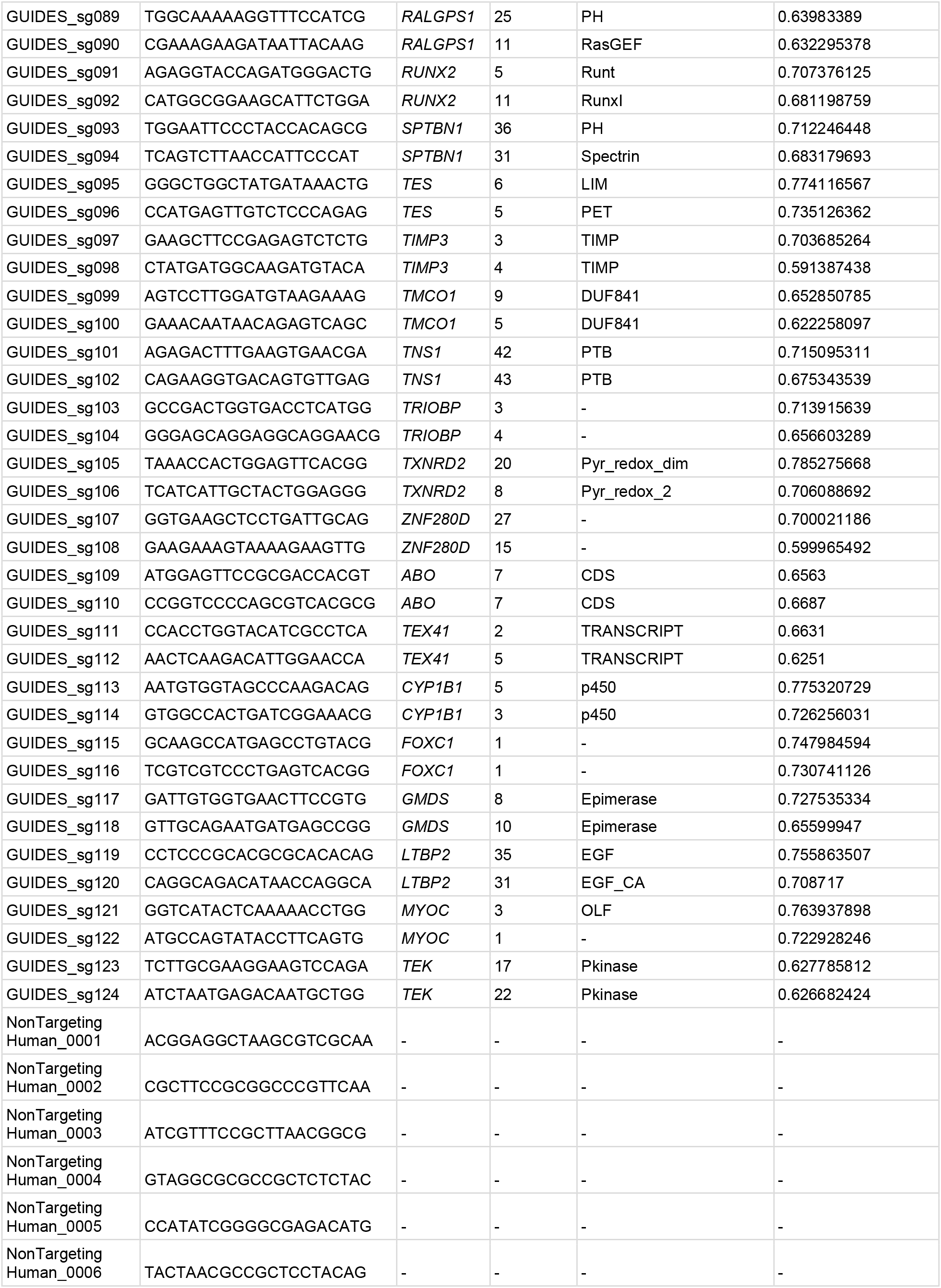
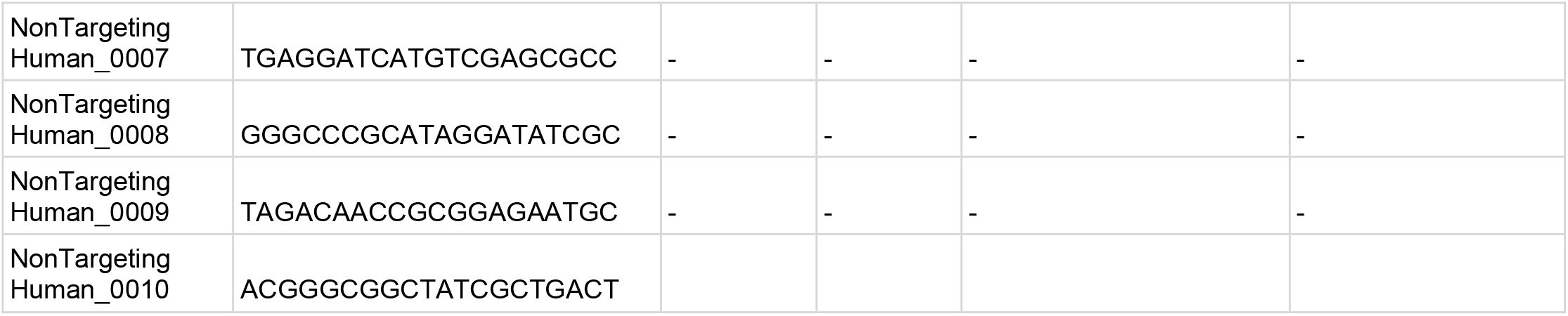

**Supplementary table 2.**
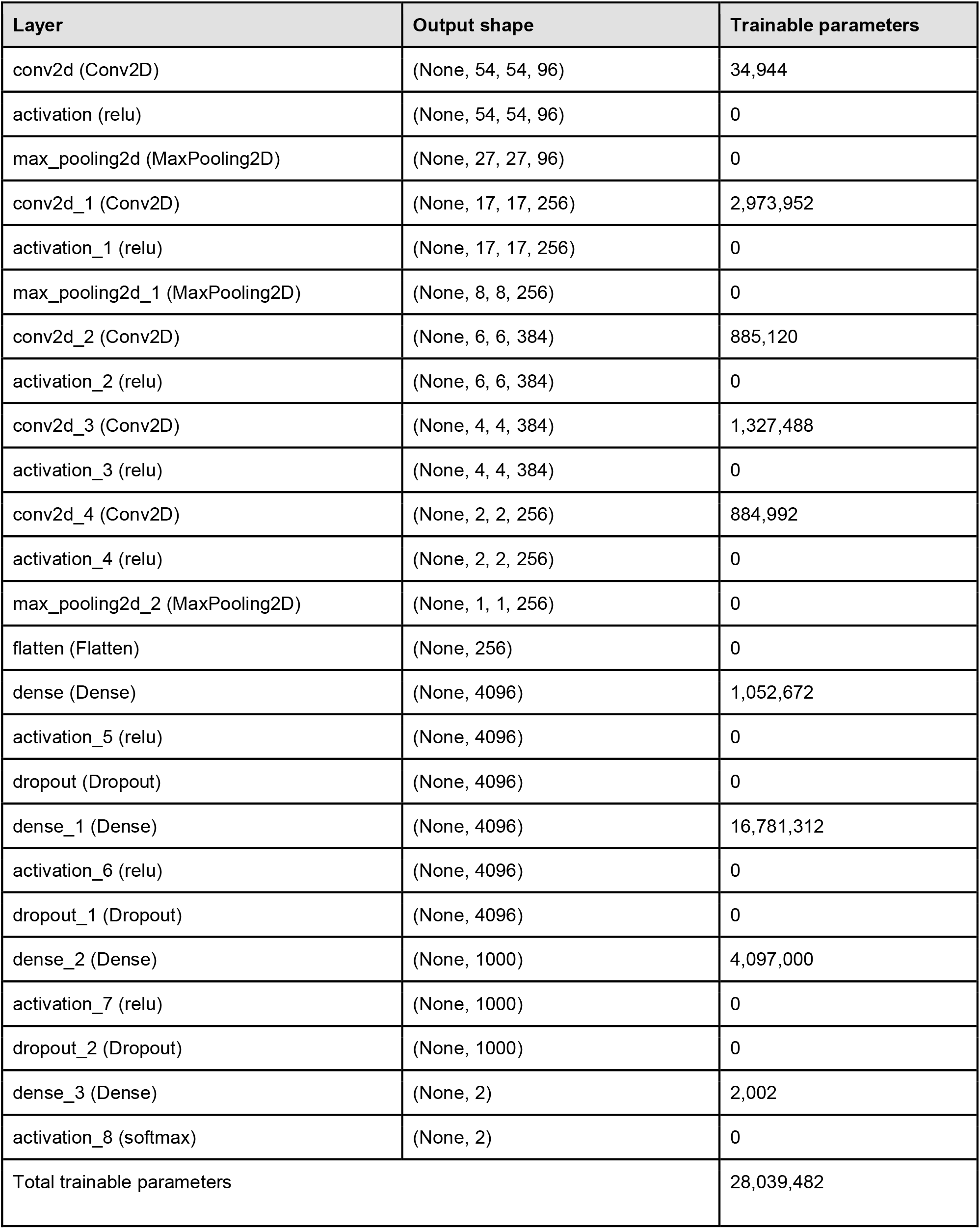
Tensorflow CNN architecture.

